# Long-Term Functional Rescue of Trauma-Induced Vision Loss by a Novel, Small Molecule TrkB Activator

**DOI:** 10.1101/2025.02.18.638863

**Authors:** Shweta Modgil, Christopher L. Walker, Micah A. Chrenek, Hans E. Grossniklaus, Frank E. McDonald, P. Michael Iuvone

## Abstract

Brain-derived neurotrophic factor (BDNF) signaling through the tropomyosin-related kinase B (TrkB) receptor promotes neuronal growth and survival following an injury. However, its short half-life and pleiotropic effects limit the clinical use of BDNF as a therapy in neurodegenerative disorders. Identification of novel and selective TrkB activators may ameliorate the damage caused to retinal neurons during eye-related injuries, and may reduce adverse visual outcomes associated with visual trauma. We previously described a selective TrkB agonist, N-[2-(5-hydroxy-1H-indol-3-yl) ethyl]-2-oxopiperidine-3-carboxamide (HIOC), that reduces the decline in visual function in a mouse model of ocular trauma (1). Using the lead optimization approach, we subsequently synthesized a fluoropyridine analog of HIOC, 2-fluoro-N-(2-(5-hydroxy-1H-indol-3-yl) ethyl) nicotinamide (HIFN), which also successfully activates TrkB. HIFN is a more potent TrkB activator than the parent compound, HIOC. Further, treatment with HIFN demonstrated neuroprotection in an animal model of overpressure ocular blast injury, ameliorating blast-related visual functional decline. Mice treated with HIFN had better visual acuity, contrast sensitivity, and retinal function supported by enhanced survival of retinal ganglion cells compared to vehicle-treated animals. Moreover, HIFN exhibited better protective effects than HIOC. The therapeutic effects of HIFN were attributed to TrkB activation, as blocking the receptor with a selective receptor antagonist (ANA-12) abrogated the neuroprotection. Together, our results identify HIFN, a novel TrkB receptor activator, as a strategy for decreasing retinal degeneration and progressive vision loss associated with traumatic ocular injury. In addition, this compound may have broader applications treating other diseases with altered TrkB activity.

## Introduction

BDNF is an endogenous ligand of the TrkB receptor, which upon its binding, induces receptor dimerization and autophosphorylation of tyrosine residues in the intracellular kinase domain of the receptor (2). TrkB/BDNF through downstream signaling molecules, phosphatidylinositol 3-kinase (PI3K)/Akt, mitogen-activated protein kinase (MAPK)/Erk, or phospholipase (PLC-γ), regulates the normal functioning of neurons and promotes survival following damage to neurons (3). TrkB/BDNF signaling has important roles in a wide range of neurodegenerative disorders ranging from Alzheimer’s and Parkinson’s diseases (4), amyotrophic lateral sclerosis, to optic neuropathies (5), as well as psychiatric disorders such as depression (6).

TrkB activation is protective in traumatic brain injury (TBI). TBI is common among civilians in blast prone-areas, and suffered by many military personnel in war zones (7). According to a report by Health.mil, 485,553 United States service members sustained at least one blast injury between 2000 and 2023Q2 (https://health.mil/Military-Health-Topics/Centers-of-Excellence/Traumatic-Brain-Injury-Center-of-Excellence/DOD-TBI-Worldwide-Numbers). With better-designed protective gear, the mortality in such cases has decreased, but the morbidity associated with these blasts is much more prevalent. Fall-related injuries in the elderly, automobile accidents, and sports injuries are also significant causes of traumatic injuries. These injuries are often accompanied by visual impairment; 75% of traumatic brain injury (TBI) patients are reported to have vision-related problems (8). Traumatic injury can include rupture of the eye globe or penetration by foreign objects resulting in an open eye injury. However, in closed globe injuries, the corneoscleral junction often remains intact, but superficial or intraocular injury is present. These injuries are more difficult to address because they usually go unnoticed, manifesting slowly without any visible symptoms. By the time they are detected, vision loss is often irreversible. Trauma-induced visual deficits may arise due to loss of retinal ganglion cells (RGCs) and optic nerve degeneration that disrupts the signal transduction to the higher brain centers. At present, there are no effective treatments for such traumatic optic neuropathy. While TrkB activation is protective in traumatic brain injury (TBI) (9–11), few studies focus on vision.

Although therapeutic effects of BDNF are evident from preclinical studies, clinical utility is limited by the short biological half-life of BDNF and its inability to cross the blood-brain barrier (BBB) and blood-retina barrier (BRB) (12–13). This has steered the field to develop alternative molecules that specifically activate the TrkB receptor in the central nervous system upon systemic administration. In this direction, synthetic peptide mimetics of BDNF including essential sequences for TrkB interaction stimulated TrkB (14) but were susceptible to proteolytic degradation. To overcome this challenge, in the last decade research has shifted towards finding potent small molecule TrkB agonists with the desired properties of penetration to the central nervous system from the systemic circulation and more stability than BDNF. We previously reported a selective TrkB agonist N-[2-(5-hydroxy-1H-indol-3-yl) ethyl]-2-oxopiperidine-3-carboxamide (HIOC, **1**, Fig 1), which passes the blood-retinal barrier (BRB) and blood-brain barrier (BBB), has a relatively long half-life *in vivo*, and mitigates visual function decline (1). In the present study, using HIOC as a lead compound, we synthesized various analogs and tested them for TrkB activation. Our data identify HIFN (**5**, **Fig 1**), an analog of HIOC, as a novel and more potent TrkB activator, and provides proof of concept for its efficacy in reducing ocular trauma-induced visual dysfunction.

**Fig 1:**
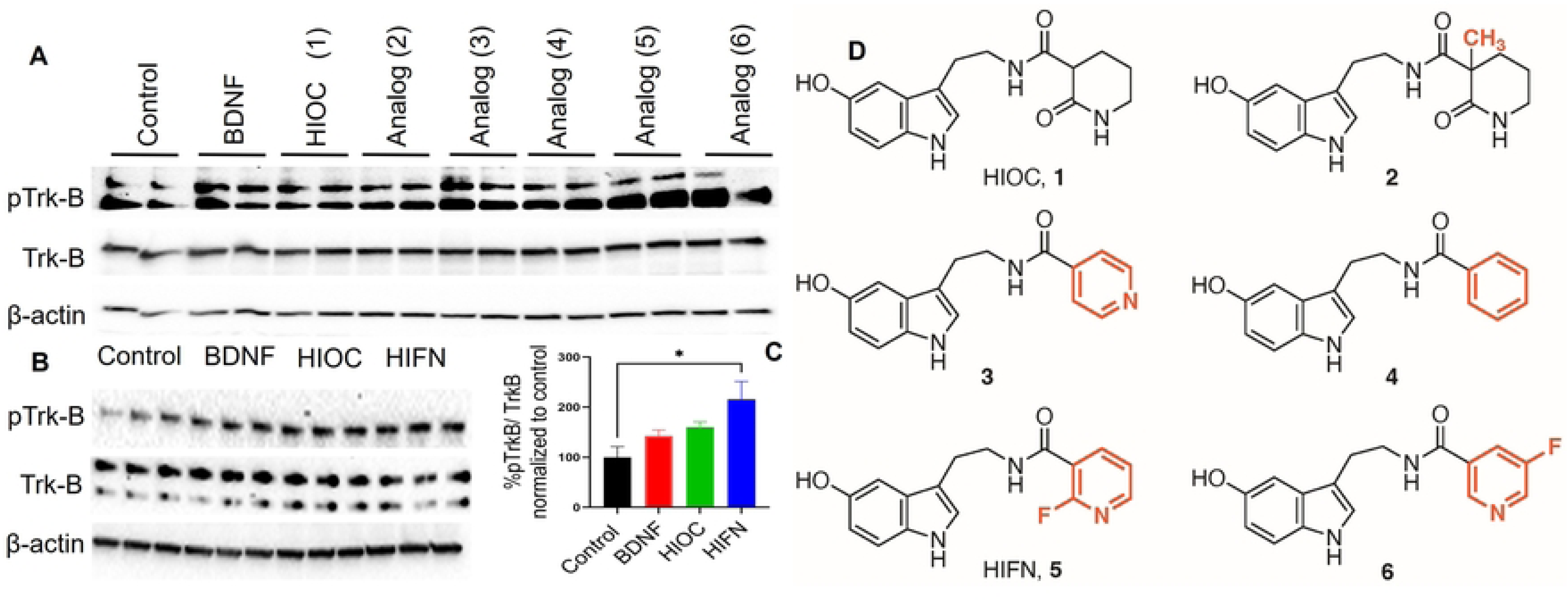
HIFN is a better TrkB activator than parent HIOC. (**A**) Cell-based screening of HIOC analogs in primary cortical neurons. Analogs were tested at 10 nM concentration and TrkB activation was assessed using western blot. Analogs **5** (HIFN) and **6** both activated TrkB and were better than HIOC. (B) TrkB activation by HIFN was also confirmed by pTrkB expression in NIH-3T3 cells expressing TrKB receptor after stimulation. (C) Quantitative analysis of TrkB phosphorylation; levels of phosphorylated TrkB were normalized to total TrkB and expressed as percentage to control (control cells were treated with DMSO, which was used to dissolve analogs). * p≤0.05 vs Control. (D) Chemical structure of parent molecule HIOC (**1**) and analogs of HIOC tested in-vitro.

## Materials and Methods

### Analog Synthesis

The synthesis of HIOC analogs generally followed the *N*-acylation protocol developed for the synthesis of HIOC (**1**) (15)., from serotonin hydrochloride and the corresponding carboxylic acid (Fig S1-S6). These optimized conditions favored selective acylation of the primary amine of serotonin hydrochloride via *N*-acylimidazole intermediates.

### Primary culture and NIH-3T3-TrkB cell line culture

Timed pregnant Sprague Dawley rats were procured from Charles River Laboratories. Primary cortical culture from embryonic day 18 rats were prepared as previously described by Pacifici and Peruzzi (16).

NIH-3T3 cells expressing human TrkB (NIH-3T3-TrkB) were provided by Frank Longo, Stanford University, and were propagated in DMEM medium (ATCC-30-2002) supplemented with 10% calf serum (ATCC-30-2030), penicillin/streptomycin (100 I.U./mL penicillin and 100 μg/mL streptomycin, P4333, Sigma-Aldrich, St Louis, MO). For protein recovery experiments, 3*10^5^ cells were plated on 6 well poly-D-lysine-coated plates (Corning) and protein extraction was carried out using cell lysis buffer (Invitrogen #FNN0011, Carlsbad, CA) containing protease inhibitor (Roche #11836153001, Basil, Switzerland) and phosphatase inhibitor (Roche, #04906845001).

### Stimulation with Analogs

Cells were serum deprived overnight to avoid the possible exposure of cells to BDNF present in serum and were stimulated with various analogs at 10 nM concentration. BDNF (#SRP3014-10UG, Sigma, 4 ng/ml) was used as a positive control for TrkB phosphorylation. Thirty minutes after the addition of test compounds, cells were washed with ice-cold PBS and treated with ice-cold lysis buffer prepared as described above. Cell lysate was collected and centrifuged at 10,000 rpm for 10 minutes @ 4°C and stored at –20°C until further use.

### Western Blotting

Total protein estimation in cell lysates was done using BCA (#23227, Thermo-Scientific). Protein lysate (50 µg) was prepared for each sample using a 4X Laemelli buffer (#1610747, Bio-Rad). Proteins were separated on 7.5-15% gradient polyacrylamide gels (#5671085, Bio-Rad) using tris-glycine-SDS buffer (#1610732, Bio-Rad) followed by transfer to 0.45 mm PVDF membranes (Midi PVDF Transfer Packs, #1704157, Bio-Rad) for antibodies staining. The membrane was blocked in 5% bovine serum albumin (BSA) (#37520, Thermo-Scientific) in Tris-buffered saline-0.1% Tween 20 (TBST) for 30 minutes at room temperature. Primary antibody incubation was carried out [Anti-phospho TrkB (#4619, Cell Signaling), anti-phospho Akt (#4060, Cell Signaling), anti-TrkB (#92991, Cell Signaling), anti-Akt (C67E7, Cell Signaling) and anti-β-actin (#4970, Cell Signaling)] overnight at 4°C, followed by washing next day. Blots were then incubated with horse radish peroxidase (HRP)-conjugated secondary antibody (#7074S, Cell Signaling) for 1 h at RT, developed with HRP substrate (#WBLUR0500, Millipore), and imaged using a Chemidoc (Bio-Rad).

### Animal and Ethics Statement

C57BL/6J mice, 2-3 months old, were procured from Jackson laboratories (Bar Harbor, ME, USA). All animal experiments were conducted in accordance with guidelines of the Care and Use of Laboratory Animals from the National Institutes of Health. All procedures were approved by Emory University’s Institutional Animal Care and Use Committee and the US Army Animal Care and Use Review Office. Mice were maintained in a standard 12:12 h light/dark cycle with free excess to food and water. Animals were anesthetized with intraperitoneal (ip) injection of a xylazine (100 mg/kg)/ketamine (10 mg/kg) cocktail, where indicated. Mice were euthanized by asphyxiation with carbon dioxide from a bottled source followed by cervical dislocation.

### Experimental Design

Animals were subjected to a blast of air directed at the front of the right eye, as described earlier (17). Briefly, the mouse was anesthetized with a mixture of xylazine/ketamine and positioned in a plexiglass sleigh that slides into a plastic polyvinyl chloride cylinder. This hollow cylinder had a hole equal to the diameter of muzzle of the air gun. The position of the mouse was adjusted in a way that the right eye of the mouse is in line with the hole and head is supported with padding to prevent secondary injuries. The cylinder provides protection to the rest of the body except the eye. The pressure was calibrated with a pressure transducer (Honeywell SensorTec sensor, Model STJE, 0-100 psi) at the muzzle of the gun and a blast of compressed air (∼20 psi at the level of the eye) was delivered. The eye was kept moist with Optixcare eye gel (Aventix, Irvine, CA) to prevent corneal drying and desiccation. After induction of blast, animals were placed on a heating pad and allowed to recover from anesthesia aided by administration of atipamezole (0.5 mg/kg ip). Sham animals were treated identically except for exposure to air pressure waves.

### Drug Preparation and Treatment

HIOC (**1**) was synthesized as described in Setterholm et al. (2015). HIFN (analog **5**) and analog **6** were synthesized (Fig S1). Stock solutions of HIOC and HIFN were prepared in 100% DMSO and further diluted with final solution containing 10% DMSO, 16.5% Cremophor EL (Sigma-Aldrich St. Louis, MO), and 16.5% ethanol in PBS. Stock solutions of ANA-12 (Sigma-Aldrich St. Louis, MO) were prepared using the same formulation as for analogs. For *in vitro* experiments, 10 nM of analogs were added to the culture medium. For animal studies, unless noted otherwise, mice were injected intraperitoneally (ip) with either analog (40 mg/kg) or vehicle, 30 min after blast and then daily at approximately the same time of day for the next 6 days (7 doses in total). For the ANA-12 experiments, the mice were pretreated (0.5 mg/kg, ip) 2.5 h prior to the daily injection of HIFN.

### Contrast Sensitivity and Visual Acuity

Visual functions in animals were measured using optometry (Cerebral Mechanics. Inc., Lethbridge, Alberta, Canada). Contrast sensitivity was measured at a spatial frequency of 0.064 cycles/degree, which is the optimal spatial frequency for C57BL/6J mice (18). Briefly, animals are kept on a raised platform inside a closed chamber surrounded by LED display screens that generate a virtual rotating drum of vertical, sinusoidal light and dark bars. The chamber is mounted with a video camera to observe the animal’s behavior in real time. Alternating light and dark vertical stripes moving in either clockwise or anticlockwise direction are presented to the mouse triggering an optomotor reflex. The contrast sensitivity threshold is the lowest contrast at which mice showed the visual reflex and data are presented as the inverse of the contrast threshold. To evaluate visual acuity, the animal was shown alternating black and white gratings (at fixed contrast of 100%) of increasing spatial frequency starting with a frequency of 0.039 c/d to find animal’s spatial frequency threshold. Contrast sensitivity follows a circadian rhythm (19); therefore, all recordings were made between 11am to 3pm. The data are presented as percent of sham control for contrast sensitivity and visual acuity.

### Scotopic electroretinography (ERG) and pattern ERG (PERG) recordings

Animals were dark adapted overnight and prepared for ERG recordings under dim red light (<3 lux). Mice were anesthetized and pupils dilated with topical application of 1% tropicamide ophthalmic solution (Akorn, Inc., Lake Forest, IL), followed by application of 1-2 drops of proparacaine hydrochloride (0.5% ophthalmic solution, Akorn, Inc.). Body temperature was maintained with a heated platform integral to the Celeris ERG system (Diagnosys, Lowel, MA). A layer of Optixcare gel was applied to cornea and stimulators were positioned with the gel making a thin layer between the cornea and stimulators. For scotopic ERG measurements, simultaneous recording of both eyes was carried out using an intensity series of white flashes ranging from 0.01 to 10 cd*s/m^2^.

PERG was recorded at 100% contrast. A stimulus of horizontal gratings with contrast reversal was used with the following specification; spatial frequency of 0.155 c/d, 2.1 contrast reversal per sec and mean luminance of 50 cd/m^2^. In total, 600 sweeps were averaged to get the final measurements. The PERG waveform is characterized by a small negative wave with an implicit time of approximately 50 msec (N1), a second wave in the form of positive deflection, peaking at approximately 80 msec (P1) followed by a late negative peak at around 350 msec (N2). PERG P1 and N2 wave amplitude ± SEM are plotted for all groups to determine functional activity of RGCs.

### Retinal wholemount staining

Animals were sacrificed, eyes were enucleated and fixed in 4% paraformaldehyde for 1 hour. The eye was dissected along the limbus, and the anterior portion containing lens and vitreous was discarded. The eyecup with retina was flattened by making radial cuts along nasal-temporal and inferior-superior retina. The retinal wholemount was washed with 1X PBS to remove any attached vitreous or iris and blocked with blocking buffer (10% donkey serum, 5% BSA, 0.5% tritonX-100 in PBS) for 1 hour at room temperature (RT). Overnight incubation at 4°C in primary antibody (anti-Brn3a; # sc-8429, Santa Cruz,) diluted in 0.5% tritonX-100/PBS (1:500) was done. Next day, following washing in 1X PBS-0.1% Tween20 (3 times), retinas were incubated in secondary antibody diluted in 1X PBS (1:1000) for 2 hours at room temperature (RT) in dark. The secondary antibody was removed, and retina was washed with PBS three times (10-15 min each, in dark). The retina was finally transferred to a glass slide and mounted in Vectashield mounting medium. Imaging was done with a Nikon Ti2 microscope with a Nikon A1R confocal imager. RGCs were counted in Image J using the RGC plug-in.

#### Statistics

Data are expressed as mean ± standard error of the mean (SEM) and were analyzed by Student’s t-test for comparisons between two groups. For multiple comparisons, one- or two-way analysis of variance as appropriate, with Tukey’s posthoc test was applied using GraphPad Prism 9. If the data failed the equal variance test, they were analyzed using the Kruskal-Wallis test.

## Results

### HIFN, a novel TrkB activator

The synthetic methods and characterization of the HIOC analogs studied is provided in Supporting Information (Fig S1-S6). We conducted cell-based screening of HIOC analogs to evaluate their efficacy in activating TrkB receptor, by employing two different cell systems (Fig 1). A fibroblast cell line (NIH-3T3-cells) stably expressing human TrkB receptor and primary cortical neurons were stimulated with various analogs for 30 minutes. Compounds **5** and **6** exhibited TrkB phosphorylation similar to BDNF (4 ng/ml), utilized as a positive control. Notably, both compounds demonstrated robust TrkB phosphorylation in primary neurons (Fig 1A). Specifically, analog **5** (2-fluoro-N-(2-(5-hydroxy-1H-indol-3-yl) ethyl) nicotinamide; hereafter, referred to as HIFN), a fluoropyridine derivative of HIOC (Fig 1D) exhibited heightened TrkB activity in NIH-3T3-TrkB cells in comparison to the parent molecule HIOC at a concentration of 10nM (Fig 1B and 1C).

### HIFN *vs* HIOC: enhanced protection in ocular trauma

In our earlier study of HIOC, we demonstrated the therapeutic potential of TrkB activators in mitigating ocular trauma-induced visual dysfunction (Dhakal et al., 2021). Prompted by our cell-based screening, we investigated the protective effects of analogs **5** and **6** against trauma injury in a mouse model of blast overpressure injury^15^. Surprisingly, analog **6**, irrespective of *in vitro* TrkB activation, showed no statistically significant protection against vision loss in blast-exposed mice (Fig S7). In contrast, analog **5** (HIFN) effectively mitigated damage caused by overpressure injury as described below.

For *in vivo* studies, animals were randomly assigned to Sham-Vehicle (Veh), Blast-Veh, Blast-HIOC, and Blast-HIFN groups. Contrast sensitivity and visual acuity recordings, a week before blast, revealed no significant intergroup baseline differences. Animals were administered a dose of 40 mg/kg intraperitoneally (ip) of HIFN or HIOC, unless specified otherwise. Treatment was initiated on the day of blast exposure and continued daily for one week (Fig. 2A).

**Fig 2.**
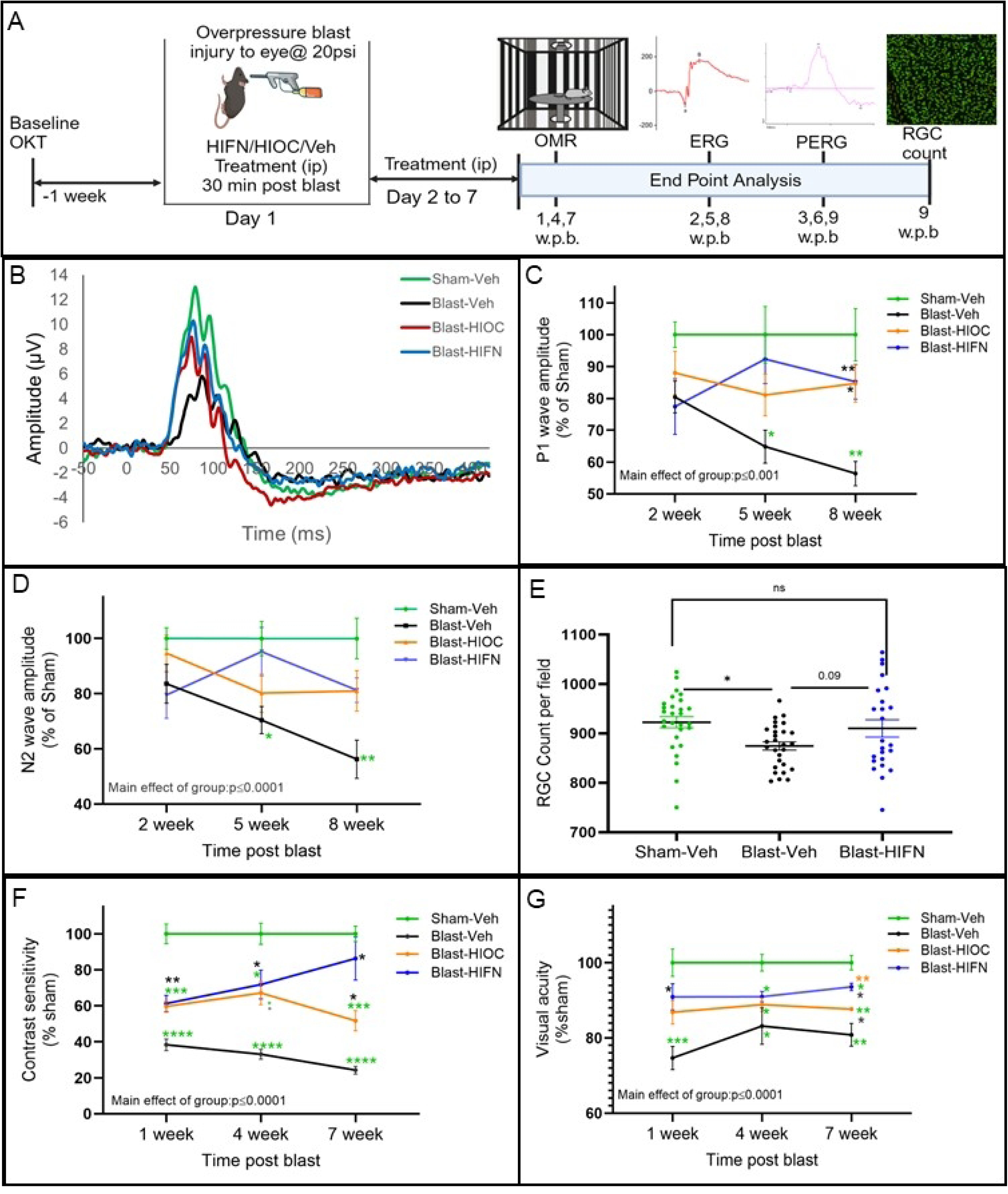
HIFN reduces retinal and visual function deficits following ocular trauma. (A) Schematic of experimental timeline. Baseline optomotor responses (OMR) were recorded in animals one week before trauma induction. The animals were randomly divided into various groups. On day 1, overpressure injury was induced by a 20 psi air blast using a modified paintball gun; 30 minutes after injury, animals were administered the first dose of vehicle, HIFN (40mg/kg ip) or HIOC (40mg/kg ip). The animals were treated for another 6 days with daily ip injections. OMR, scotopic electroretinogram (ERG) and pattern electroretinogram (PERG) were recorded at different timepoints as indicated in figure; week post blast (w.p.b). (B) Representative PERG waveform from each group. (C and D) Pattern ERG recorded for retinal ganglion cell function assessment at various times post blast. P1 (C) and N2 (D) wave amplitudes begin to decline progressively over the 8 weeks. HIFN treatment prevented the decline in RGC cell function and preserved the P1 and N2 amplitude at 8-week post blast. E) Retinal wholemounts were immunostained with RGC specific marker Brn3a. Cells in ROI were automatically counted using ImageJ, Simple RGC plugin. RGC cell count decreases after blast in vehicle-treated mice while HIFN treatment prevented the RGC cell loss. Contrast sensitivity (F) and visual acuity (G) at 1,4 and 7-week post blast. Data is presented as % of Sham-Veh ± SEM. HIFN rescued visual acuity and contrast sensitivity deficit at all the time points. At 7-weeks visual function is better in HIFN treated animals compared to HIOC. (C) and (D) * p≤0.05 vs Sham-Veh, ** p≤0.01 vs Sham-Veh; * p≤0.05 vs Blast-Veh; ** p≤0.01 vs Blast Veh; n=5-7per group. Data are expressed as %Sham-Veh ± SEM; REML, Holm Sidak’s adjusted p-values used for post-hoc comparisons. (E) Data are expressed as total Brn3a-positive cells/ROI (mean ± SEM) * p≤0.05; n=5-8/group. (F) and (G) * p≤0.05 vs Sham-Veh, ** p≤0.01 vs Sham-Veh, ***p≤0.001 vs Sham-Veh, ****p≤0.0001 vs Sham-Veh; * p≤0.05 vs Blast-Veh, ** p≤0.01 vs Blast-Veh; ** p≤0.01vs Blast-HIOC; n=6-7per group; REML, Holm Sidak’s adjusted p-values used for post-hoc comparisons.

To understand which retinal cell types are affected by blast injury, we recorded electroretinograms (ERG) to map the functional activity of retinal cells. Full-field scotopic ERG recordings (–3.0 to 1 log cd.s/m^2^) at 3, 6 and 9 weeks after blast showed no observable difference for any ERG component across time (Table S2; Fig S8). No significant differences were observed in photoreceptor and bipolar cell function as analyzed by a-wave (main group effect, F (3, 110) = 1.1.3, p=0.340) and b-wave amplitudes (main group effect, F (3,110) = 0.848, p=0.470). We further isolated oscillatory potentials OPs and c-wave amplitudes from ERGs at 9-weeks. OPs arise from amacrine cell responses on the ascending limb of the ‘b’-wave while the c-wave indicates retinal pigment epithelial cell function. We observed no alterations in ERG c-wave amplitudes in any treatment group (p=0.244; Fig S8-C) suggesting normal functioning of retinal pigment epithelial cells. Similarly, the amplitudes of OPs were unaltered after the blast exposure (p=0.117; Fig S8-D). These findings indicated that ocular trauma under the conditions used in our study did not significantly impair photoreceptor, bipolar cell, and amacrine cell light responses.

Traumatic injuries often affect retinal ganglion cells given the vulnerability of their axons as they extend long distances to brain centers (Moham et al., 2013). Since the function of outer or inner retinal neurons appeared intact, we next examined the functionality of retinal ganglion cells (RGCs) using the pattern ERG (PERG). PERG revealed significant changes in both P1 and N2 wave amplitudes [2-way, mixed model ANOVA, {P1 amplitude main group effect F (3,21)=11.21, p=0.0001}; {N2 amplitude, main group effect F (3,81) =11.80, p=0.0001}]. In blast-exposed animals treated with vehicle, both P1 and N2 amplitudes progressively declined over 8 weeks (Fig 2B,C,D). Initial assessment 2 weeks after blast did not show any significant differences between blast-exposed and sham animals. However, 5 weeks after blast, both P1 (p=0.03, vs Sham-Veh) and N2 wave amplitudes (p=0.017, vs Sham-Veh) decreased in vehicle-treated animals. RGC function appeared better in animals with HIFN treatment as reflected by P1 amplitude, but the effect was not statistically significant (p= 0.07 vs Blast-Veh). A further assessment of RGC function at 8 weeks post-blast confirmed a continued decline in the vehicle treatment group (p=0.007 for P1 amplitude; p=0.005 for N2 amplitude vs Sham-Veh) suggesting a slow but progressive decline in RGC cell function after blast. At this time point both HIFN and HIOC rescued the deficit in RGC cell function as evidenced by significantly higher P1 wave amplitudes in HIOC (p=0.01 vs Blast-Veh) and HIFN (p=0.007 vs Blast-Veh) treated mice. Moreover, N2 wave amplitude modestly improved in drug-treated animals (p=0.051, HIFN vs Veh).

To correlate the RGC function with the cell number, we prepared retinal whole-mounts from animals sacrificed after PERG recordings and stained them for Brn3a, a cell-specific marker for RGCs. Regions of interest (ROI) were selected from the central (600 μm from the optic disc) region of the retina. Quantification of Brn3a+ cells revealed a small loss of RGCs in the retina of Blast-Veh mice compared to Sham (p=0.019; Fig 2E). Blast-HIFN mice showed no significant decline in RGCs compared to Sham, and a trend for an increased number of RGCs compared to Blast-Veh mice (p=0.09).

Blast exposure significantly impacted visual function outcomes, assessed by the optomotor response (OMR), with a decline in contrast sensitivity evident as early as 1 week after blast exposure, when it was reduced to 38.4 ±3.2% (p=0.0001) of sham control animals (Fig 2F). Contrast sensitivity continued to decline (24.3 ± 2.1%, p=0.0001) with time until 7-weeks post-blast (longest time point recorded). Preservation of contrast sensitivity with either HIOC (59.6 ± 3.2%, p=0.002 vs Veh) or HIFN (61.3± 4.5%, p=0.007vs Veh) was observed at 1-week post-blast. By 4 weeks, contrast sensitivity declined by 67% in vehicle group while decline in animals treated with HIOC (33%, p=0.01 vs Veh) and HIFN (39%, p=0.01 vs Veh) was significantly less. Notably, at 7 weeks after blast, contrast sensitivity of HIFN-treated mice was 3-fold higher than that of vehicle-treated mice and was not significantly different than that of sham controls (p=0.32) (Fig 2F). Moreover, HIFN-treated animals demonstrated a 22.3% higher (p=0.06 vs HIOC) contrast sensitivity than the HIOC group at 7 weeks after blast, which was similar to sham group (p=0.32 vs Sham).

Visual acuity deficits, as assessed by spatial frequency threshold in OMR, began at 1-week post-blast (Fig 2G) in parallel to lower sensitivity to contrast, although compared to contrast sensitivity, visual acuity showed less sensitivity to blast. Visual acuity showed a 25.2% decline one-week post-blast (p= 0.001 vs sham). HIOC and HIFN treatment resulted in better visual acuity preservation compared to the vehicle group. 7 weeks after blast, the visual acuity was 8.3% higher with HIOC (p=0.06) and 16% with HIFN (p=0.01) compared to vehicle-treated animals. Furthermore, HIFN was significantly better than HIOC in reducing the visual acuity deficit (p=0.002). Together, these results indicate that BDNF mimetics can prevent the functional decline in retinal cell function and provide better vision outcomes after ocular injury. Moreover, HIFN exhibited long-lasting protective effects, superior to the parent molecule HIOC in preventing vision loss due to traumatic injury.

The comprehensive study aimed to evaluate the protective effects and efficacy of HIFN was initially performed on male C57BL6/J mice. In order to determine if these effects were sex-specific, trauma was induced in female C57BL6/J mice as well. Visual acuity deficits were recorded in both HIFN-treated and non-treated males and females 8 days after blast. The treatment with HIFN reduced the visual acuity decline as shown (Fig S9) in both sexes to a comparable degree.

### Neuroprotection by HIFN is TrkB dependent

HIFN potently activated the TrkB receptor (Fig 1) suggesting that the preservation of RGC and visual function is conferred via TrkB mediated signaling. To test this hypothesis, on day 1, 2.5 h before the blast, animals received a systemic injection of the TrkB receptor-specific inhibitor, ANA-12 (0.5mg/kg)^25^ or its vehicle. Subsequently, 30 minutes after blast, animals were treated with HIFN or vehicle. Additionally, from day 2 to 7, ANA-12/vehicle was injected 2.5h before HIFN/vehicle treatment to interfere with the TrkB activation (Fig 3A). The study aimed to assess the impact of ANA-12 pretreatment on the neuroprotective effects exerted by HIFN.

**Fig 3.**
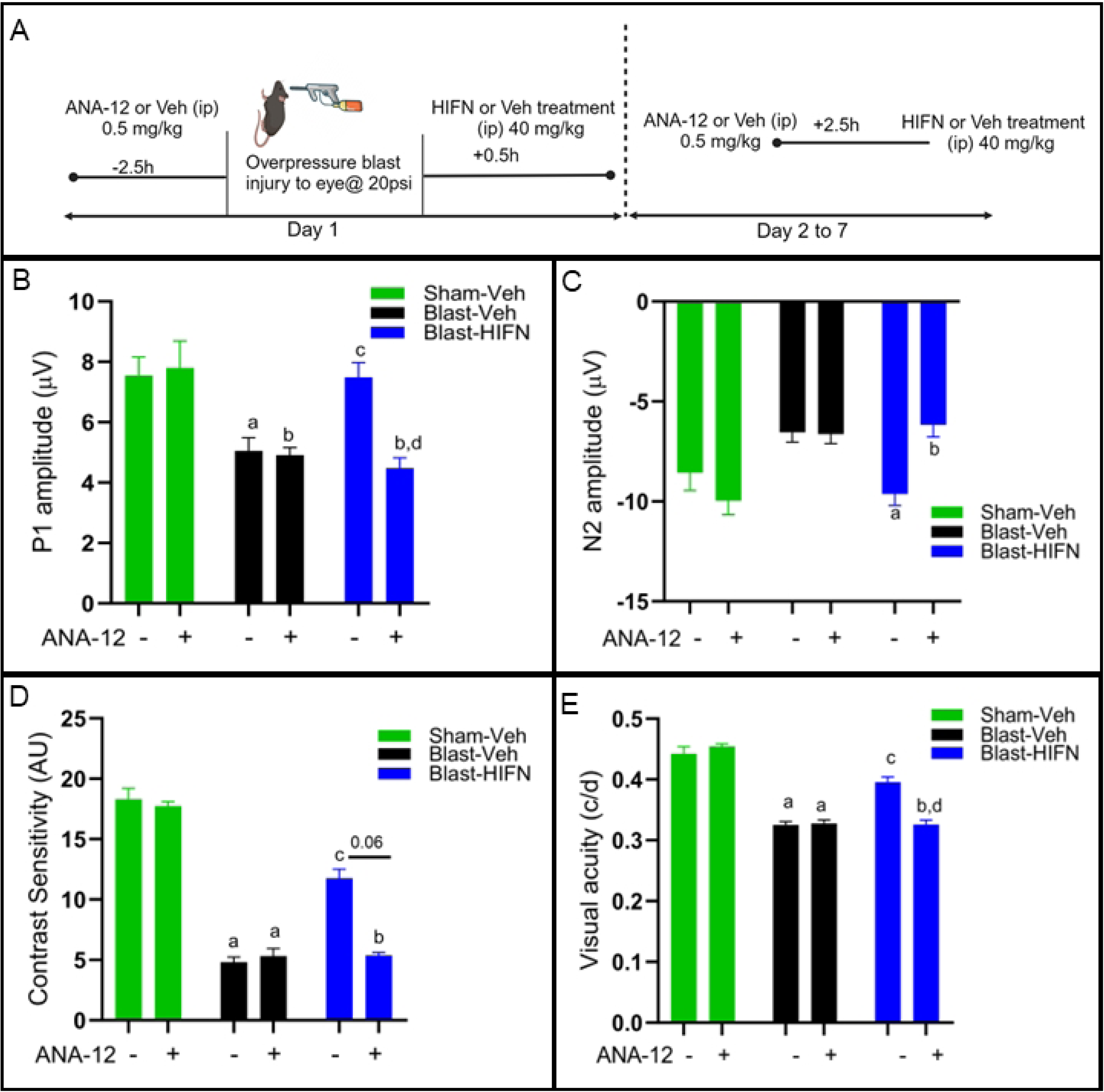
HIFN protection of vision loss is Trk-B dependent: (**A**) Schematic representation of experimental protocol of ANA-12 and HIFN treatment. ANA-12, a specific TrkB receptor inhibitor, was used to block TrkB activity. On day 1, animals received 0.5 mg/kg of either ANA-12 or vehicle 2.5 h prior to blast injury. Animals were administered HIFN or vehicle 30 min after blast exposure. ANA-12 pretreatment followed by vehicle/HIFN was continued for a week. (B) and (C) PERG was recorded at 8 w.p.b.; preservation of P1 and N2 wave amplitude by HIFN was not seen in animals that were pretreated with ANA-12. (D) and (E) Contrast sensitivity and visual acuity were recorded at 7 w.p.b.; Contrast sensitivity and visual acuity decline in HIFN treated animals were seen when animals were prior treated with ANA-12. (A) **^a^**p≤0.05 vs Veh/Sham-Veh; **^b^**p≤0.01 vs Veh/Sham-Veh; **^c^**p≤0.05 vs Veh/Blast-Veh; **^d^**p≤0.01vs Veh/Blast-HIFN (B) **^a^**p≤0.05 vs Veh/Blast-Veh; **^b^**p≤0.01vs Veh/Blast-HIFN; n=7-9/group (C) **^a^**p≤0.0001 vs Veh/Sham-Veh; **^b^**p≤0.001 vs Veh/Sham-Veh; **^c^**p≤0.05 vs Veh/Blast-Veh (D) **^a^**p≤0.0001 vs Veh/Sham-Veh; **^b^**p≤0.001 vs Veh/Sham-Veh; **^c^**p≤0.05 vs Veh/Blast-Veh; **^d^**p≤0.05 vs Veh/Blast-HIFN

As discussed above, HIFN reduced the loss of RGC function induced by traumatic injury (Fig 2B-2D). Pretreatment with ANA-12 eliminated these protective effects of HIFN on RGC function (Fig 3B,3C). ANA-12 alone had no significant effect on the P1 amplitude of the PERG in the Sham mice (p=0.99) or on the reduction in amplitude following blast (p=0.99) (Fig 3B). However, ANA-12 completely blocked the protective effect of HIFN administration; the P1 amplitude in the ANA-12 pretreatment group (ANA-12/Blast-HIFN) was significantly lower (p=0.003) compared to the group not treated with ANA-12 prior to HIFN (Veh/Blast-HIFN), and not significantly different from the ANA-12-treated group not administered HIFN. Likewise, N2 amplitude showed a similar pattern (Fig 3C) where ANA-12 had no effect alone (Sham, p=0.65; Blast p=0.97), but blocked the increased amplitude caused by HIFN (p=0.008).

Additionally, ANA-12 blocked visual function improvements with HIFN (Fig 3D and 3E) in mice exposed to blast injury. Similar to RGC function, ANA-12 alone had no effect on contrast sensitivity or visual acuity in Sham or blast exposed mice. HIFN reduced the contrast sensitivity deficit in mice not pretreated with ANA-12 (Veh/Blast vs HIFN/Blast, p=0.02), but not in those pretreated with ANA-12. Contrast sensitivity in mice treated with ANA-12 and HIFN was comparable to that in mice treated with ANA-12 alone (p=0.61). Visual acuity threshold revealed the same pattern as contrast sensitivity. Animals treated with HIFN following blast exposure have significantly (p=0.017) better visual acuity threshold than vehicle treated mice. The improvement in visual acuity with HIFN treatment was not seen in ANA-12/Blast-HIFN animals. The mice treated with ANA-12 and HIFN had visual acuity similar to the mice treated with ANA-12 alone and significantly lower (p=0.026) than the HIFN alone group. Taken together, the data showed that pharmacological inhibition of TrkB activation blocked the protection exerted by HIFN, supporting our hypothesis that HIFN acts via TrkB receptor mediated signaling.

### Neuroprotective effects of HIFN are dose dependent

All our experiments were conducted with 40 mg/kg of HIFN, which was the optimal dose found for HIOC in our previous study (Dhakal et al., 2021). Aiming to identify the optimal dosage for achieving neuroprotection with HIFN, we next conducted a dose-response study. HIFN was injected into animals at doses ranging from 1 to 100 mg/kg. A 30 mg/kg dose significantly reduced the decline in P1 (vs Blast-Veh p=0.013) and N2 (Blast-Veh, p=0.004) wave amplitudes 8 weeks after blast exposure (Fig 4A and 4B). Animals treated with 1 mg/kg, 3 mg/kg, and 10 mg/kg HIFN following blast-exposure showed diminished RGC function similar to vehicle-treated animals, indicating that these lower doses were ineffective in rescuing RGC damage. The highest dose tested (100 mg/kg) had significantly improved N2 amplitude (vs Blast-Veh, p=0.04), but improvement in P1 amplitude, though better than vehicle treatment, did not reach statistical significance (p=0.32).

**Fig 4.**
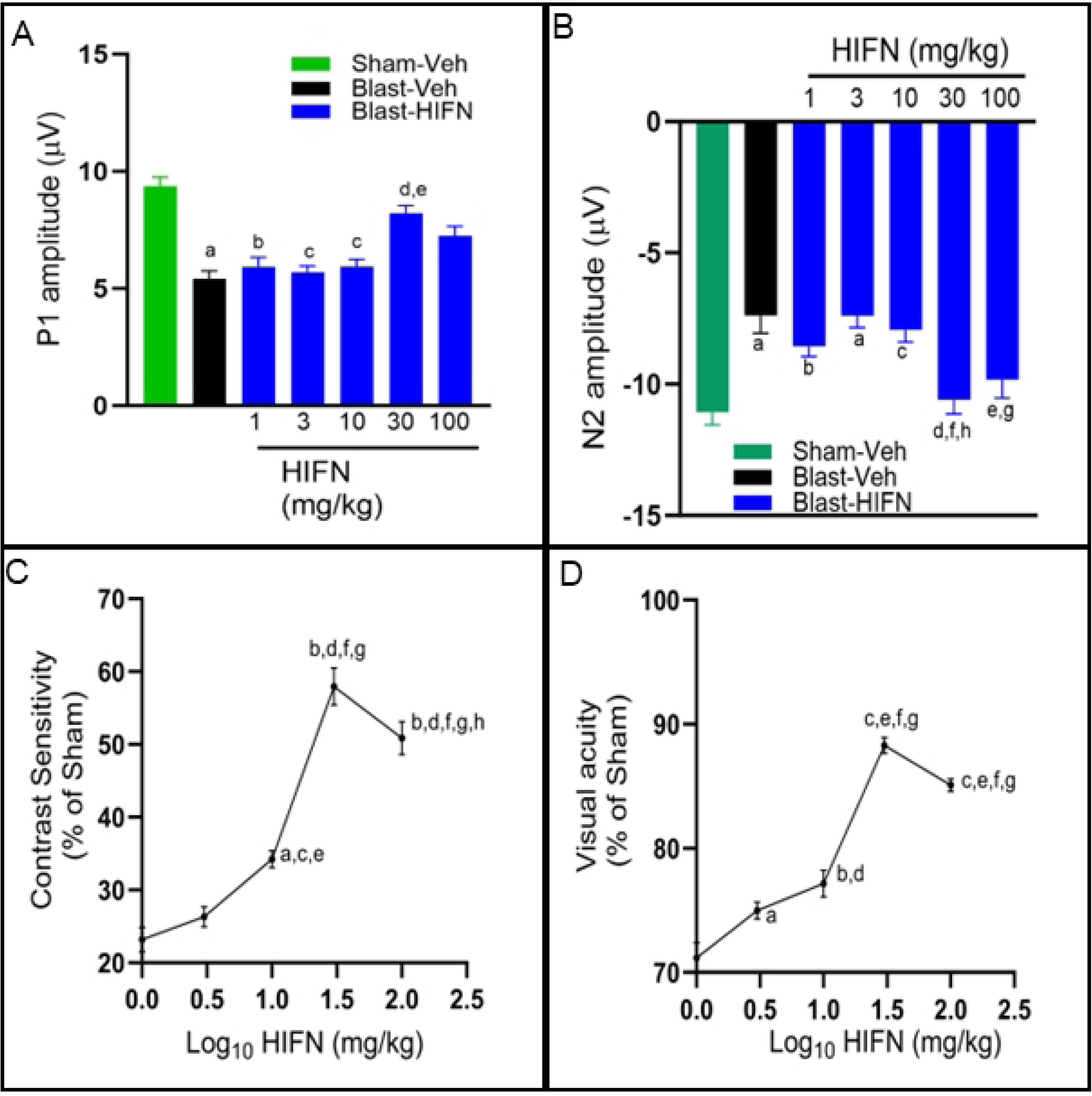
Dose dependent neuroprotection by HIFN. Various doses of HIFN were tested to find the effective dose. The best responses were observed with 30 mg/kg for reducing RGC and visual function dysfunctions. (A) **^a^**p≤0.001 vs Sham-Veh; **^b^**p≤0.05 vs Sham-Veh; **^c^**p≤0.01 vs Sham-Veh; **^d^**p≤0.05 vs Blast-Veh; **^e^**p≤0.05 vs 3mg/kg (B) **^a^**p≤0.001 vs Sham-Veh; **^b^**p≤0.05 vs Sham-Veh; **^c^**p≤0.01 vs Sham-Veh; **^d^**p≤0.01 vs Blast-Veh; **^e^**p≤0.05 vs Blast-Veh; **^f^**p≤ 0.01vs3mg/kg; **^g^**p≤0.05 vs 3mg/kg; **^h^**p≤0.05 vs 10mg/kg; n= 6-8/group (C) ^a^p≤0.01 vs vehicle; **^b^**p≤0.0001vs vehicle; ^c^p≤0.01 vs 1 mg/kg; **^d^**p≤0.001 vs 1mg/kg; **^e^**p≤0.05 vs 3mg/kg; **^f^**p≤0.0001 vs 3mg/kg; **^g^**p= p≤0.0001 vs 10mg/kg; **^h^**p≤0.05 vs 30mg/kg (D) **^a^**p≤0.05 vs vehicle; **^b^**p≤0.001vs vehicle; **^c^**p≤0.0001 vs vehicle; **^d^**p≤0.001 vs 1mg/kg; **^e^**p≤0.0001 vs 1mg/kg; **^f^**p≤0.0001 vs 3mg/kg; **^g^**p≤0.0001 vs 10mg/kg

We recorded dose-dependent preservation of visual function 7 weeks after blast exposure, with an apparent peak dose of 30 mg/kg (Fig 4C,4D). We saw significant improvement in contrast sensitivity (Fig 4C) starting from a dose of 10 mg/kg (vs Blast-Veh, p=0.008) and continuing to improve with higher doses of 30 mg/kg and 100 mg/kg (vs Blast-10mg/kg, p=0.0001). Likewise, visual acuity increased significantly from 71.2± 1.2% of sham control to 85.1± 0.5% (p=0.0001) with HIFN dose increasing from 1 mg/kg to 100 mg/kg HIFN (Fig 4D). An HIFN dose as low as 3 mg/kg was effective in visual acuity preservation compared to vehicle (p=0.019) (Fig 4D). Maximum visual acuity preservation (88.3 ± 0.64 % of sham control) was observed with the 30 mg/kg dose (vs Blast-Veh, p=0.0001). Overall, these results show a dose dependent neuroprotection by HIFN. Acuity and contrast preservation were achieved starting with a dose of 10 mg/kg, but RGC function improvement required a higher dose. In summary, in our experiments, animals treated with 30 mg/kg had the best visual outcomes.

### Therapeutic time window for HIFN-induced neuroprotection is limited to 3 h after injury

To determine the therapeutic treatment window for HIFN, we injected the initial dose of HIFN at different time intervals after blast. Mice received 40 mg/kg dose of HIFN starting at 0.25, 1, 3, 6 or 24 h after blast exposure; mice then received daily injections at the same time of day for an additional 6 days. We recorded changes in visual function and RGC function at 7-weeks and 8-weeks post blast, respectively. HIFN treatment mitigated vision loss if provided within 3 h of injury (Fig 5). Beyond the 3 h time-window, HIFN loses its efficacy in rescuing the vision decline.

**Fig 5.**
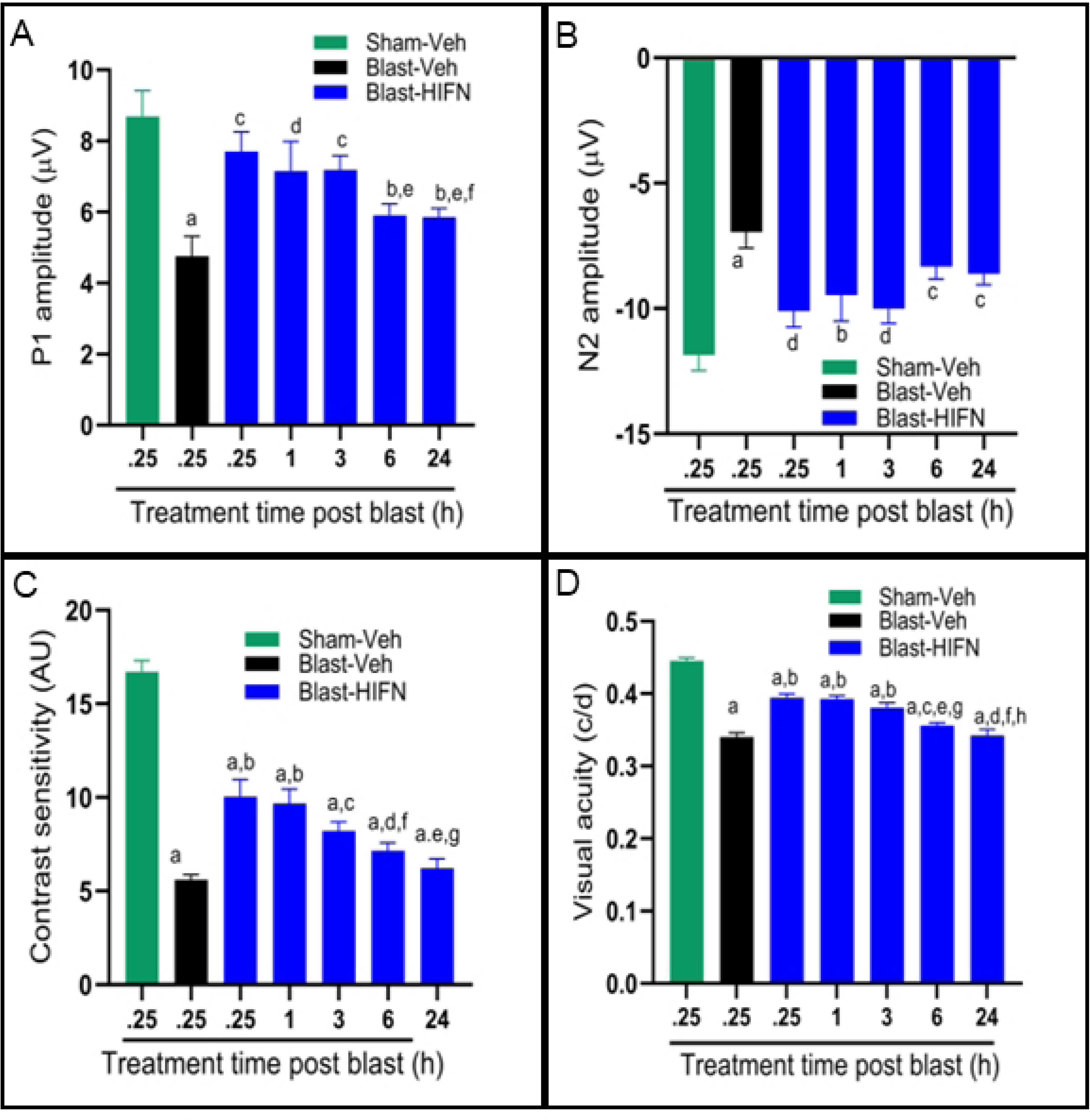
Critical time window for HIFN intervention: Mice received their initial treatment with HIFN (40mg/kg) at various times relative to blast exposure (treatment time post blast) and were treated daily for the next 6 days. Initial treatment was provided as early as 15 minutes (0.25 h) after the blast and delayed up to 24 h. PERG P1-wave amplitude (A) and N2-wave amplitude (B), contrast sensitivity (C), and visual acuity (D) were recorded. HIFN reduced vision loss with initial treatments up to 3 h, but was less effective at 6 h or 24 h. A) **^a^**p≤0.0001 vs Sham-Veh; **^b^**p≤0.01 vs Sham-Veh; **^c^**p≤0.01 vs Blast-Veh; ^d^p≤0.05 vs Blast-Veh; ^e^p≤0.05 vs 0.25h; **^f^**p≤ 0.05 vs 3h; B) **^a^**p≤0.0001 vs Sham-Veh; **^b^**p≤0.05 vs Sham-Veh; ^c^p≤0.01 vs Sham-Veh; ^d^p≤0.01 vs Blast-Veh; n= 6-8/group. C) **^a^**p≤0.0001 vs Sham-Veh; **^b^**p≤0.0001 vs Blast-Veh; **^c^**p≤0.05 vs Blast-Veh; mg/kg; ^d^p≤0.05 vs 0.25 h; **^e^**p≤0.001 vs 0.25h; **^f^**p≤ 0.05 vs 1h; **^g^**p≤0.01 vs 1h; D) **^a^**p≤0.0001 vs Sham-Veh; **^b^**p≤0.0001 vs Blast-Veh; **^c^**p≤0.001 vs 0.25h; **^d^**p≤0.0001 vs 0.25h; **^e^**p≤0.001 vs 1h; ^f^p≤ 0.0001 vs 1h; **^g^**p≤0.05 vs 3h; **^h^**p≤0.001 vs 3h; n=6-8/group

There was improved function of RGC cells as recorded by PERG in animals treated with HIFN within a time window of 3 h after injury. P1 amplitude (p=0.001, p=0.02, p=0.002 for 0.25 h, 1 h and 3 h, respectively, vs Blast-Veh) and N2 amplitude (p=0.004, p=0.06, p=0.002 for 0.25 h, 1 h and 3 h, respectively, vs Blast-Veh; Figure 5A and 5B) were significantly higher in animals treated with HIFN. RGC function was not rescued if the initial HIFN treatment was delayed for 6 hours or more. Visual function data supported PERG results. HIFN treatment preserved contrast sensitivity even if the treatment was delayed up to 3 h (vs Blast-Veh, p=0.014). However, providing treatment within 1 h gave the best response. Contrast sensitivity was 60.7 ± 5.5% (p=0.0001) for 0.25 h and 57.9±4.6%, (p=0.0001) for the 1 h group compared to 33.6 ± 1.57% for vehicle treated animals (Fig 5C and 5D). A treatment window not exceeding 3 h reduced visual acuity deficits. Visual acuity was significantly better than Blast-Veh group in animals treated at 0.25 h (p=0.0001), 1 h (p=0.0001) and 3 h (p=0.0001) after blast. If the first dose of HIFN was administered 6 or 24 h after blast exposure, the visual acuity deficit was not significantly reduced.

### Promising Safety Profile of HIFN in Toxicity Testing

Safety profile of HIFN was assessed using acute and chronic toxicity studies in both male and female animals. For acute toxicity studies, animals were subjected to a dose of 300 mg/kg and monitored for toxicity including change in fur color, lethargy, red secretions around eye, hunch back, tremors, as well as mortality for 7-day period. Remarkably, at this dose, no mortality or observable signs of toxicity were noted at this dose. Subsequently, a higher dose of 600 mg/kg was administered to another cohort of animals. Even at this elevated level, 20 times the maximal effective dose for visual function protection, all animals remained active, displaying no visible signs of toxicity.

For the evaluation of chronic toxicity, male and female mice received a daily dose of 40 mg/kg HIFN for a duration of 40 days. Body weights of vehicle and HIFN-treated mice were not significantly different at baseline or after treatment for 40 days (baseline males: Veh 30.0 ± 0.6 g, HIFN 29.8 ±0.6 g; baseline females: Veh 18.8 ± 0.3 g, HIFN 18.8 ± 0.6 g; post-treatment males: Veh 29.5 ± 0.5 g, HIFN 29.7 ± 0.3 g; post-treatment females: Veh 19.2 ± 0.3 g, HIFN 18.6 ± 0.4 g). Blood and serum samples were collected for complete blood count and serum biochemistry analyses. Brain, liver, kidney, spleen, and heart were harvested for histopathological examination of sections. In male mice, we observed no significant differences in blood parameters or liver and kidney function tests between the control and HIFN-treated groups (Table 1 and Table 2). HIFN-treated females exhibited an increase in the number of white blood cells and lymphocytes; however, aside from these changes, all other parameters remained comparable to the control group. Importantly, there were no evident signs of histopathological toxicity upon analysis of stained tissue sections (Fig 6). In conclusion, the results from both chronic and acute toxicity studies indicate that HIFN is relatively non-toxic. The absence of mortality or observable signs of toxicity at high doses in acute toxicity assessments, coupled with the lack of significant adverse effects on various physiological parameters and organ histopathology in chronic toxicity studies, supports the safety profile of HIFN. These findings collectively suggest that HIFN holds promise as a safe therapeutic agent.

**Table 1:**
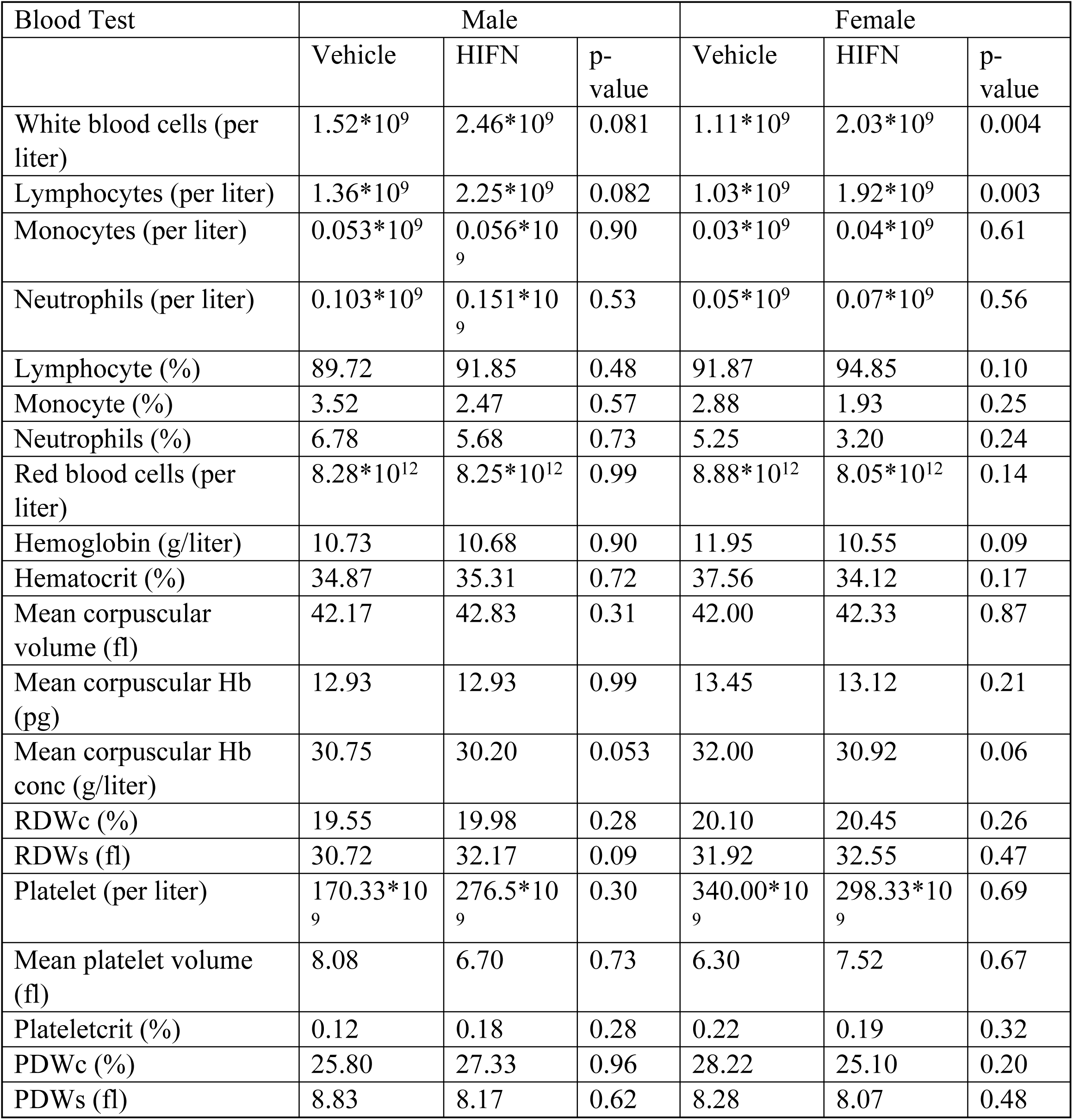
CBC profile of HIFN and vehicle treated male and female animals. n=6 animals/group Abbreviations: RDWc: RBC distribution width, coefficient of variation; RDWs: RBC distribution width, standard deviation; PDWc: Platelet distribution width, coefficient of variation; PDWs: Platelet distribution width, standard deviation.

| Blood Test | Male |  |  | Female |  |  |
| --- | --- | --- | --- | --- | --- | --- |
|  | Vehicle | HIFN | p-value | Vehicle | HIFN | p-value |
| White blood cells (per liter) | 1.52*10<sup>9</sup> | 2.46*10<sup>9</sup> | 0.081 | 1.11*10<sup>9</sup> | 2.03*10<sup>9</sup> | 0.004 |
| Lymphocytes (per liter) | 1.36*10<sup>9</sup> | 2.25*10<sup>9</sup> | 0.082 | 1.03*10<sup>9</sup> | 1.92*10<sup>9</sup> | 0.003 |
| Monocytes (per liter) | 0.053*10<sup>9</sup> | 0.056*10<sup>9</sup> | 0.90 | 0.03*10<sup>9</sup> | 0.04*10<sup>9</sup> | 0.61 |
| Neutrophils (per liter) | 0.103*10<sup>9</sup> | 0.151*10<sup>9</sup> | 0.53 | 0.05*10<sup>9</sup> | 0.07*10<sup>9</sup> | 0.56 |
| Lymphocyte (%) | 89.72 | 91.85 | 0.48 | 91.87 | 94.85 | 0.10 |
| Monocyte (%) | 3.52 | 2.47 | 0.57 | 2.88 | 1.93 | 0.25 |
| Neutrophils (%) | 6.78 | 5.68 | 0.73 | 5.25 | 3.20 | 0.24 |
| Red blood cells (per liter) | 8.28*10<sup>12</sup> | 8.25*10<sup>12</sup> | 0.99 | 8.88*10<sup>12</sup> | 8.05*10<sup>12</sup> | 0.14 |
| Hemoglobin (g/liter) | 10.73 | 10.68 | 0.90 | 11.95 | 10.55 | 0.09 |
| Hematocrit (%) | 34.87 | 35.31 | 0.72 | 37.56 | 34.12 | 0.17 |
| Mean corpuscular volume (fl) | 42.17 | 42.83 | 0.31 | 42.00 | 42.33 | 0.87 |
| Mean corpuscular Hb (pg) | 12.93 | 12.93 | 0.99 | 13.45 | 13.12 | 0.21 |
| Mean corpuscular Hb conc (g/liter) | 30.75 | 30.20 | 0.053 | 32.00 | 30.92 | 0.06 |
| RDWc (%) | 19.55 | 19.98 | 0.28 | 20.10 | 20.45 | 0.26 |
| RDWs (fl) | 30.72 | 32.17 | 0.09 | 31.92 | 32.55 | 0.47 |
| Platelet (per liter) | 170.33*10<sup>9</sup> | 276.5*10<sup>9</sup> | 0.30 | 340.00*10<sup>9</sup> | 298.33*10<sup>9</sup> | 0.69 |
| Mean platelet volume (fl) | 8.08 | 6.70 | 0.73 | 6.30 | 7.52 | 0.67 |
| Plateletcrit (%) | 0.12 | 0.18 | 0.28 | 0.22 | 0.19 | 0.32 |
| PDWc (%) | 25.80 | 27.33 | 0.96 | 28.22 | 25.10 | 0.20 |
| PDWs (fl) | 8.83 | 8.17 | 0.62 | 8.28 | 8.07 | 0.48 |

**Fig 6.**
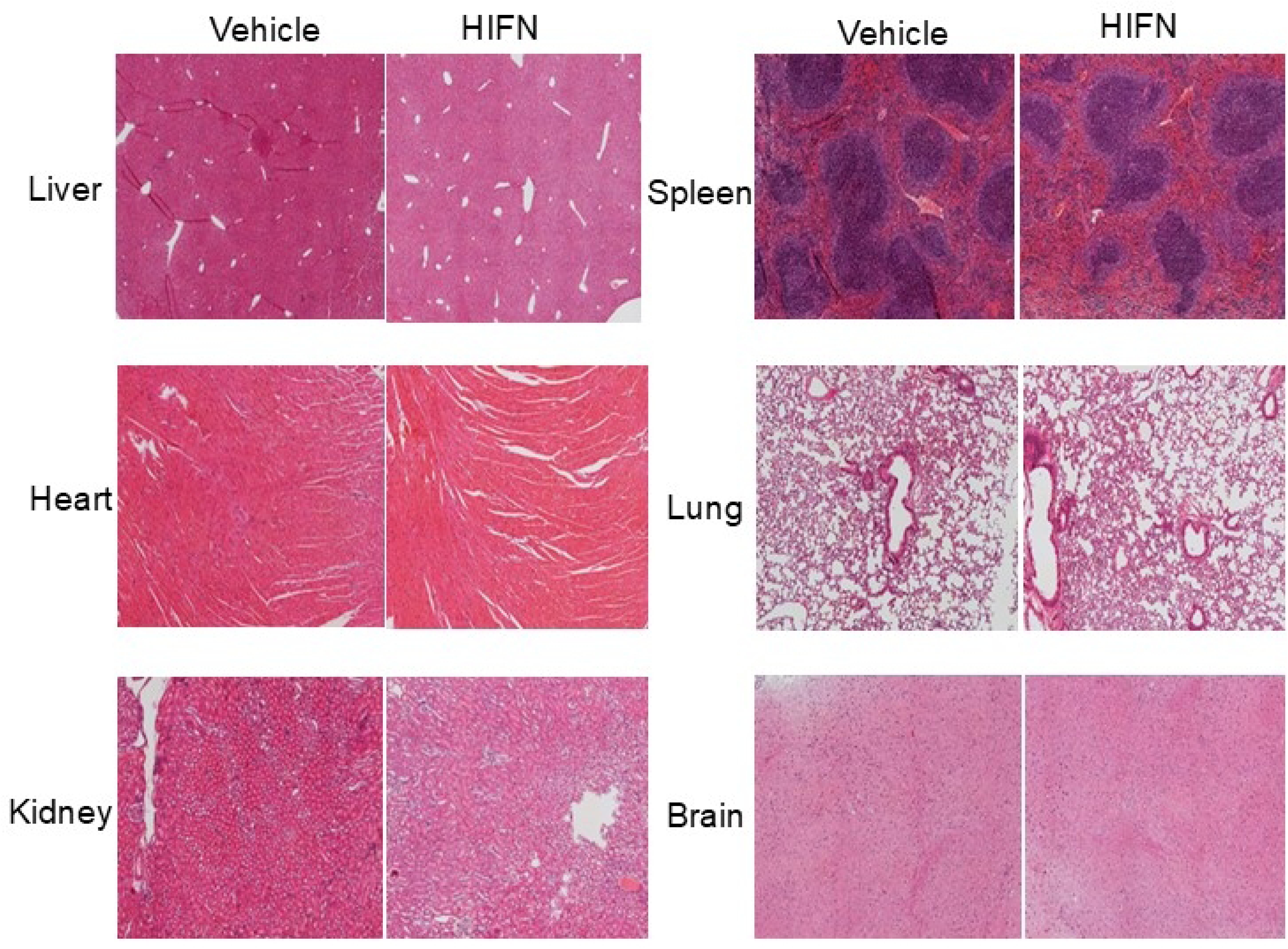
No signs of toxicity of HIFN treatment were reported in histopathological analysis. Representative images of hematoxylin/eosin-stained tissue sections from animals treated with HIFN or vehicle for 40 days. The sections were analyzed by a board-certified pathologist (HEG). No abnormal changes due to HIFN were observed. n=3/group

**Table 2:** Serum chemistry to evaluate kidney and liver function in the animals after chronic HIFN treatment. n= 6 animals/group Abbreviations: ALP: Alkaline phosphatase, ALT: alanine transaminase; AMYL: Amylase; AST: aspartate transaminase; BA-SEK: blood albumin:, BUN: blood urea nitrogen; CREAT: creatinine; DBILI: direct bilirubin, LDH: lactate dehydrogenase; TBILI: total bilirubin

| Test | Male |  |  | Female |  |  |
| --- | --- | --- | --- | --- | --- | --- |
|  | Vehicle | HIFN | p-value | Vehicle | HIFN | p-value |
| ALB (g/dL) | 2.44±0.24 | 2.56±0.074 | 0.80 | 2.7±0.08 | 2.55±0.09 | 0.33 |
| ALP (U/L) | 37.6±6.46 | 39.6±1.32 | 0.297 | 70.3±5.49 | 64.16±2.53 | 0.332 |
| ALT (U/L) | 47.6±8.84 | 58.8±9.47 | 0.412 | 64.33±18.51 | 61.66±4.93 | 0.89 |
| AMYL (U/L) | 836.8±224.44 | 668.8±24.72 | 0.478 | 624.667±28.09 | 553.5±42.85 | 0.08 |
| AST (U/L) | 533.2±126.67 | 473.2±80.59 | 0.690 | 467.33±131.09 | 510±89.60 | 0.79 |
| BA-SEK (µmol/L) | 19.9±9.92 | 8.82±1.00 | 0.30 | 32.45±20.27 | 22.72±7.83 | 0.818 |
| BUN (mg/dl) | 29.6±3.96 | 26±3.63 | 0.53 | 22.33±1.58 | 19.5±1.087 | 0.339 |
| CREAT (mg/dl) | 0.272±0.05 | 0.424±0.03 | 0.051 | 0.35±0.04 | 0.43±0.04 | 0.206 |
| DBILI (mg/dl) | 0.44±0.16 | 0.44±0.07 | 0.99 | 0.36±0.08 | 0.35±0.06 | 0.99 |
| LDH (U/L) | 1414.8±317.82 | 1258.8±198.41 | 0.68 | 1037.33±294.24 | 1215.83±193.63 | 0.62 |
| TBILI (mg/dl) | 0.556±0.116 | 0.476±0.05 | 0.54 | 0.413±0.06 | 0.49±0.07 | 0.461 |

## Discussion

Lack of effective treatment for traumatic eye injuries has encouraged extensive research into the mechanisms underlying progressive damage resulting from these injuries. Numerous studies provide evidence supporting the involvement of RGC death (20) and TrkB signaling in blast-related injuries (21–23). A positive correlation is established between BDNF/TrkB levels and functional outcomes post-injury. However, the pharmacokinetic constraints of BDNF limit its therapeutic use. Recently, nanoparticle-bound BDNF, that could readily cross the blood-brain barrier, demonstrated effectiveness in treating traumatic blast injury (24), highlighting the importance of TrkB signaling in such injuries. Recognizing the potential of TrkB agonists, synthetic small molecules such as 7,8-dihydroxyflavone (DHF) and LM22A4 have shown promise in attenuating injury-related pathology (14). Our previous study showcased the efficacy of a small molecule, HIOC (**1**), a synthetic derivative of N-acetylserotonin, in preventing blast-induced vision loss in mice^1^. Here, we introduce HIFN (**5**) demonstrating improved efficacy over HIOC in diminishing progressive vision loss following ocular trauma. HIFN is an achiral analog of chiral HIOC having a fluoropyridine group in place of the six-membered lactam ring of HIOC. Individual enantiomers of HIOC are not configurationally stable, due to the position of the chiral center between two carbonyl groups, which promotes facile enolization (Figure S1-S2). Although each enantiomer of HIOC may exhibit distinct levels or types of bioactivities, the achiral nature of HIFN provides another advantage for HIFN over HIOC as a therapeutic agent.

A decline in contrast sensitivity and visual acuity is an established marker of traumatic eye injury (17, 25). Our study, utilizing the same overpressure blast model, corroborated these findings by demonstrating a continuous decline in spatial frequency thresholds and contrast threshold in blast-exposed animals (Figure 2). Both HIOC and HIFN preserved the contrast sensitivity and visual acuity in animals for at least two-months after the blast. Notably, the protection afforded by HIFN surpassed that of HIOC, making it a promising candidate for further investigations.

Pattern ERG (PERG), a functional readout of retinal ganglion cells, revealed that blast injury primarily affected RGCs (Figures 2,3). Our results align with other studies reporting alterations in RGC function following injury (26). The retina is a complex structure wherein intermediatory bipolar cells pass on the signal generated in photoreceptors in the outer retinal layers to the retinal ganglion cells. Any damage to outer or inner neurons also disrupts signaling to RGC and affects their responses. Unlike some studies reporting subnormal (27) and supernormal ERG responses (25) post-injury, our scotopic ERG data did not reveal involvement of outer or inner nuclear layer neurons. These results were consistent with our previous study where amplitudes of ERG a- and b-waves were unaltered after the blast injury (1). This discrepancy could be attributed to several procedural factors including distance of the eye from the blast source, pressure of blast wave, the type of driver gas used (helium, nitrogen, etc.), and animal species/strain (28). In our model, blast overpressure injury significantly decreased the RGC cell function without affecting ERG a- or b-waves. The preservation of PERG function with HIFN suggests a preventive effect on RGC dysfunction. Considering the large decrements of contrast sensitivity and reversal by HIFN, the changes in RGC cell number were remarkably small. The reason for this apparent discrepancy is unclear, but may involve changes in the optic nerve such as disruptions of myelination, that did not cause RGC cell death. Further studies involving optic nerve recording or analyzing cell death using apoptotic markers may provide valuable insight about mechanisms of vision preservation by HIFN.

Our investigation into the underlying neuroprotective mechanism of HIFN revealed its dependence on TrkB receptor activation. The administration of a specific TrkB receptor blocker, ANA-12 (29), negated the protective effect of HIFN on vision loss. This was not surprising, as HIOC, the parent analog of HIFN, also exerts its protection through TrkB signaling (1,30). Although HIFN was given for only the first week after blast exposure, *the protection was sustained or improved over the subsequent two months*. TrkB/BDNF signaling is involved in neuronal survival and plasticity. We propose that HIFN has promoted neuronal survival and aided surviving neurons in making new connections, by activating TrkB.

The preservation of visual function was dose dependent, with an apparent maximal response at 30 mg/kg. Interestingly, the highest tested dose (100 mg/kg) showed a marginal decline in protective effects, potentially hinting at off-target binding issues at higher concentrations. Off-target binding may stimulate other unknown pathways. The critical therapeutic window in response to traumatic blast injury for HIFN was identified as 3 hours, beyond which the neuroprotective effects were not realized. A plausible reason is that irreversible damage may have occurred at latencies beyond this time.

Small molecule TrkB activators like HIFN offer promise for neurodegenerative disorders where BNDF/TrkB signaling is compromised. The present study not only confirms HIFN as a TrkB receptor activator but also provides evidence of its efficacy in preserving vision decline resulting from traumatic injuries. The exact mechanism by which a small molecule, particularly one with a molecular weight less than 500 Daltons, fits into the active site of the TrkB dimer and triggers its activation remains unknown. Current hypotheses suggest that small molecules like HIFN may interact with a TrkB monomer, inducing structural changes in the receptor that facilitate dimer formation^2^. Alternatively, HIFN may act on preformed TrkB dimers (31). In this scenario, small molecules could bind to these preexisting dimers, altering their conformation and promoting phosphorylation, which is a key step in receptor activation. The intricacies of how these small synthetic molecules precisely bind to and activate the TrkB receptor are areas of ongoing research. Future studies will likely delve into the details of the molecular interactions, the specific binding sites involved, and the conformational changes induced by these small molecules in the context of TrkB receptor activation. Understanding these mechanisms will not only contribute to our knowledge of neuroprotection but may also pave the way for the development of targeted therapies for neurodegenerative disorders.

In summary, HIFN is a promising TrkB receptor activator, as a potential alternative to BDNF therapy in the realm of treating neurodegenerative diseases. The demonstrated efficacy of HIFN in preserving vision and preventing progressive damage following traumatic injury suggests its potential utility in addressing a wide range of neurological and psychiatric disorders characterized by compromised BDNF/TrkB signaling.

## Acknowledgements

This study was supported by the Department of Defense, through the US Army Medical Research and Development Command under Awards W81XWH-18-1-0700, (PMI,FEM), by the NIH P30EY06360 (Emory Vision Core), by a Challenge Grant from Research to Prevent Blindness, and by a gift from the Abraham J. and Phyllis Katz Foundation. The authors thank Frank Longo and Tao Yang, Stanford University, for providing NIH-3T3-TrkB cells. We thank Dr. John Bacsa, Emory University X-ray Crystallography Center, for obtaining crystal structures of HIOC (**1**) and HIFN (**5**). Tiffany Hung, Emory University, assisted with gram-scale synthesis of HIFN (**5**). We thank Jendayi Dixon for technical assistance. Opinions, interpretations, conclusions, and recommendations are those of the authors and are not necessarily endorsed by the Department of Defense or other funding agencies.

## Author contributions

S.M., F.E.M., P.M.I. designed research; S.M., C.L.W., M.A.C. performed research; H.E.G. assessed histopathology; S.M., F.E.M., P.M.I. wrote the paper; all authors read and edited the manuscript.

## Data Availability

Crystallographic data have been deposited in the Cambridge Crystallographic Data Centre (CCDC) deposition numbers 2366286 (HIFN) and 2366626 (HIOC). All other data are included in the article and/or supporting information.

## Supporting Information

**Figure S1-1**. Chemical synthesis of HIOC (**1**) and analogs **2** - **6**. Differences in analog structures from HIOC are highlighted in red.

**Figure S2.** Time course of protium-deuterium exchange with HIOC (**1**) at methine carbon, at pD 7.8

**Figure S3**: Thermal ellipsoid representation of the asymmetric unit of HIOC (**1**). There is one molecule of the target compound in the asymmetric unit, which is represented by the reported sum formula. In other words: Z is 2 and Z’ is 1. The chiral centers have both R and S-configuration in this crystal.

**Figure S4**: Plot of a portion of the hydrogen bonding network in the crystal structure of HIOC (**1**).

**Figure S5**: The molecular structure of HIFN (**5**).

**Figure S6**: Hydrogen bonded dimer in the crystal structure of HIFN (**5**).

**Figure S7. Analog 6 did not exhibit in-vivo neuroprotection.** Animals were treated with analog **6** (ip, 40mg/kg) following blast (same regime was followed as for HIFN (**5**) treatment shown in Figure 2A). No statistically significant change in P1 (A) and N2 (B) wave was seen in animals treated with analog **6** compared to vehicle treated animals. Analog **6** treatment did not rescue the visual function deficit; no statistically significant improvement in contrast sensitivity (C) and visual acuity (D) was observed in treated animals. * p≤0.05; **p≤0.01 ;****p≤0.0001, n=4-5/group.

**Figure S8. Outer retinal neurons were not affected by overpressure blast injury.** At 9 weeks post blast, animals did not show any change in rod photoreceptor or rod bipolar cell function as assessed by ERG: (A) amplitude of ‘a-wave’; (B) amplitude of ‘b-wave’. Function of RPE and amacrine cells was also not altered as shown by: (C) ‘c-wave’ and (D) oscillatory potentials (OP), respectively. No significant differences were found for any of the parameters recorded. Data are expressed as mean ± SEM; n=6-8/group.

**Figure S9. HIFN reduces visual function decline in females and males.** Neuroprotection by HIFN was tested in female mice. Mice were exposed to blast and treated with HIFN (40 mg/kg) or vehicle 30 minutes later and daily for the next 6 days. Visual acuity was measured 8 days after blast exposure. **p≤0.01; ***p≤0.001; ****p≤0.0001, n=5-6/group.

**Table S1.** Tabulated ^13^C and ^1^H NMR data for HIFN (**5**)

**Table S2.** Electroretinogram ‘a-wave’ and ‘b-wave’ recordings 3 and 6-week post blast showing no significant effect on rod photoreceptor and bipolar cell function.

## References

1. Dhakal S, He L, Lyuboslavsky P, Sidhu C, Chrenek MA, Sellers JT, Boatright JH, Geisert EE, Setterholm NA, McDonald FE, Iuvone PM. A Tropomycin-Related Kinase B Receptor Activator for the Management of Ocular Blast-Induced Vision Loss. Journal of Neurotrauma. 2021 Oct 15;38(20):2896–906.

2. Longo FM, Massa SM. Small-molecule modulation of neurotrophin receptors: a strategy for the treatment of neurological disease. Nature reviews Drug discovery. 2013 Jul;12(7):507–25.

3. Wang Y, Liang J, Xu B, Yang J, Wu Z, Cheng L. TrkB/BDNF signaling pathway and its small molecular agonists in CNS injury. Life sciences. 2024 Jan 1;336:122282.

4. Zuccato C, Cattaneo E. Brain-derived neurotrophic factor in neurodegenerative diseases. Nature Reviews Neurology. 2009 Jun;5(6):311–22.

5. Bikbova G, Oshitari T, Baba T, Yamamoto S. Neurotrophic factors for retinal ganglion cell neuropathy-with a special reference to diabetic neuropathy in the retina. Current Diabetes Reviews. 2014 May 1;10(3):166–76.

6. Wang CS, Kavalali ET, Monteggia LM. BDNF signaling in context: From synaptic regulation to psychiatric disorders. Cell. 2022 Jan 6;185(1):62–76.

7. Dengler BA, Agimi Y, Stout K, Caudle KL, Curley KC, Sanjakdar S, Rone M, Dacanay B, Fruendt JC, Phillips JB, Meyer AC. Epidemiology, patterns of care and outcomes of traumatic brain injury in deployed military settings: implications for future military operations. Journal of trauma and acute care surgery. 2022 Aug 1;93(2):220–8.

8. Frick KD, Singman EL. Cost of military eye injury and vision impairment related to traumatic brain injury: 2001–2017. Military medicine. 2019 May 1;184(5-6):e338–43.

9. Ding Y, Zhu W, Kong W, Li T, Zou P, Chen H. Edaravone attenuates neuronal apoptosis in hippocampus of rat traumatic brain injury model via activation of BDNF/TrkB signaling pathway. Archives of Medical Science: AMS. 2019 Nov 18;17(2):514.

10. Wu CH, Hung TH, Chen CC, Ke CH, Lee CY, Wang PY, Chen SF. Post-injury treatment with 7, 8-dihydroxyflavone, a TrkB receptor agonist, protects against experimental traumatic brain injury via PI3K/Akt signaling. PloS one. 2014 Nov 21;9(11):e113397.

11. Fletcher JL, Dill LK, Wood RJ, Wang S, Robertson K, Murray SS, Zamani A, Semple BD. Acute treatment with TrkB agonist LM22A-4 confers neuroprotection and preserves myelin integrity in a mouse model of pediatric traumatic brain injury. Experimental neurology. 2021 May 1;339:113652.

12. Wurzelmann M, Romeika J, Sun D. Therapeutic potential of brain-derived neurotrophic factor (BDNF) and a small molecular mimics of BDNF for traumatic brain injury. Neural regeneration research. 2017 Jan 1;12(1):7–12.

13. Miranda-Lourenço C, Ribeiro-Rodrigues L, Fonseca-Gomes J, Tanqueiro SR, Belo RF, Ferreira CB, Rei N, Ferreira-Manso M, de Almeida-Borlido C, Costa-Coelho T, Freitas CF. Challenges of BDNF-based therapies: From common to rare diseases. Pharmacological Research. 2020 Dec 1;162:105281.

14. Massa SM, Yang T, Xie Y, Shi J, Bilgen M, Joyce JN, Nehama D, Rajadas J, Longo FM. Small molecule BDNF mimetics activate TrkB signaling and prevent neuronal degeneration in rodents. The Journal of clinical investigation. 2010 May 3;120(5):1774–85.

15. Setterholm NA, McDonald FE, Boatright JH, Iuvone PM. Gram-scale, chemoselective synthesis of N-[2-(5-hydroxy-1H-indol-3-yl) ethyl]-2-oxopiperidine-3-carboxamide (HIOC). Tetrahedron letters. 2015 Jun 3;56(23):3413–5.

16. Pacifici M, Peruzzi F. Isolation and culture of rat embryonic neural cells: a quick protocol. Journal of visualized experiments: JoVE. 2012 May 24(63):3965.

17. Hines-Beard J, Marchetta J, Gordon S, Chaum E, Geisert EE, Rex TS. A mouse model of ocular blast injury that induces closed globe anterior and posterior pole damage. Experimental eye research. 2012 Jun 1;99:63–70.

18. Prusky GT, Douglas RM. Characterization of mouse cortical spatial vision. Vision research. 2004 Dec 1;44(28):3411–8.

19. Jackson CR, Ruan GX, Aseem F, Abey J, Gamble K, Stanwood G, Palmiter RD, Iuvone PM, McMahon DG. Retinal dopamine mediates multiple dimensions of light-adapted vision. Journal of Neuroscience. 2012 Jul 4;32(27):9359–68.

20. Mohan K, Kecova H, Hernandez-Merino E, Kardon RH, Harper MM. Retinal ganglion cell damage in an experimental rodent model of blast-mediated traumatic brain injury. Investigative ophthalmology & visual science. 2013 May 1;54(5):3440–50.

21. Rostami E, Krueger F, Plantman S, Davidsson J, Agoston D, Grafman J, Risling M. Alteration in BDNF and its receptors, full-length and truncated TrkB and p75NTR following penetrating traumatic brain injury. Brain research. 2014 Jan 13;1542:195–205.

22. Ko IG, Kim SE, Hwang L, Jin JJ, Kim CJ, Kim BK, Kim H. Late starting treadmill exercise improves spatial leaning ability through suppressing CREP/BDNF/TrkB signaling pathway following traumatic brain injury in rats. Journal of exercise rehabilitation. 2018 Jun;14(3):327.

23. Kaplan GB, Vasterling JJ, Vedak PC. Brain-derived neurotrophic factor in traumatic brain injury, post-traumatic stress disorder, and their comorbid conditions: role in pathogenesis and treatment. Behavioural pharmacology. 2010 Sep 1;21(5-6):427–37.

24. Khalin I, Alyautdin R, Wong TW, Gnanou J, Kocherga G, Kreuter J. Brain-derived neurotrophic factor delivered to the brain using poly (lactide-co-glycolide) nanoparticles improves neurological and cognitive outcome in mice with traumatic brain injury. Drug delivery. 2016 Nov 21;23(9):3520–8.

25. Allen RS, Motz CT, Feola A, Chesler KC, Haider R, Ramachandra Rao S, Skelton LA, Fliesler SJ, Pardue MT. Long-term functional and structural consequences of primary blast overpressure to the eye. Journal of neurotrauma. 2018 Sep 1;35(17):2104–16.

26. Harper MM, Gramlich OW, Elwood BW, Boehme NA, Dutca LM, Kuehn MH. Immune responses in mice after blast-mediated traumatic brain injury TBI autonomously contribute to retinal ganglion cell dysfunction and death. Experimental eye research. 2022 Dec 1;225:109272.

27. Morriss NJ, Conley GM, Hodgson N, Boucher M, Ospina-Mora S, Fagiolini M, Puder M, Mejia L, Qiu J, Meehan W, Mannix R. Visual dysfunction after repetitive mild traumatic brain injury in a mouse model and ramifications on behavioral metrics. Journal of Neurotrauma. 2021 Oct 15;38(20):2881–95.

28. Chandra N, Sundaramurthy A, Gupta RK. Validation of laboratory animal and surrogate human models in primary blast injury studies. Military medicine. 2017 Mar 1;182(suppl_1):105–13.

29. Cazorla M, Prémont J, Mann A, Girard N, Kellendonk C, Rognan D. Identification of a low–molecular weight TrkB antagonist with anxiolytic and antidepressant activity in mice. The Journal of clinical investigation. 2011 May 2;121(5):1846–57.

30. Shen J, Ghai K, Sompol P, Liu X, Cao X, Iuvone PM, Ye K. N-acetyl serotonin derivatives as potent neuroprotectants for retinas. Proceedings of the National Academy of Sciences. 2012 Feb 28;109(9):3540–5.

31. Shen J, Maruyama IN. Brain-derived neurotrophic factor receptor TrkB exists as a preformed dimer in living cells. Journal of molecular signaling. 2012 Dec;7(1):1–7.

